# Tapping into the aging brain: *In vivo* microdialysis reveals mirroring pathology between preclinical models and patients with Alzheimer’s disease

**DOI:** 10.1101/2021.01.23.427888

**Authors:** C. Bjorkli, C. Louet, T.H. Flo, M. Hemler, A. Sandvig, I. Sandvig

**Affiliations:** Department of Neuromedicine and Movement Science, Faculty of Medicine, Norwegian University of Science and Technology; Center for Molecular Inflammation Research, Faculty of Medicine, Norwegian University of Science and Technology; Department of Clinical Neuroscience, Neuro, Head and Neck, Umeå University Hospital; Department of Community Medicine and Rehabilitation, Neuro, Head and Neck, Umeå University Hospital

**Keywords:** Aβ, Tau, Biomarkers, Protein aggregation, CSF, Neurodegenerative disease, 3R

## Abstract

Preclinical models of Alzheimer’s disease (AD) can provide valuable insights into the onset and progression of the disease, such as changes in concentrations of amyloid-β (Aβ) and tau in cerebrospinal fluid (CSF). However, such models are currently underutilized due to limited advancement in techniques that allow for longitudinal monitoring of CSF akin to methods employed in AD patients. An elegant way to understand the biochemical environment in the diseased brain is intracerebral microdialysis, a method that has until now been limited to short-term observations, or snapshots, of the brain microenvironment. Our novel push-pull microdialysis method in AD mice permits *in vivo* longitudinal monitoring of dynamic changes of Aβ and tau in CSF and allows for better translational understanding of CSF biomarkers. Specifically, we demonstrate that CSF concentrations of Aβ and tau along disease progression in transgenic mice mirror what is observed in patients, with a decrease in CSF Aβ observed when plaques are deposited, and an increase in CSF tau once tau pathology is present in the brain. We found that a high molecular weight cut-off membrane allowed for simultaneous sampling of Aβ and tau, comparable to lumbar puncture CSF collection in patients. We furthermore provide specific recommendations for optimal application of our novel microdialysis method, such as achieving optimal recovery of analytes, which depends heavily on the flow rate of perfusion, probe properties and perfusate composition. Our approach can further advance AD research by following evolving neuropathology along the disease cascade via consecutive sampling from the same animal, and can additionally be used to administer pharmaceutical compounds and assess their efficacy in treating AD.

## Introduction

Preclinical models of Alzheimer’s disease (AD) can provide valuable insights that can promote our understanding of disease onset and mode of progression. Importantly, such models can have high translational potential. AD animal models, such as 3xTg AD mice^1^, harboring human genetic mutations associated with a risk for developing AD neuropathology, such as amyloid precursor protein (APP), presenilin 1 and 2 (PSEN1, 2)^2^ and microtubule-associated protein tau (MAPT)^3^ have the potential to help us understand the changing biochemical environment of the brain, something that is limited to CSF sampling distal from the brain via lumbar puncture in patients. Microdialysis is an *in vivo* bioanalytical sampling technique used to monitor the analyte concentrations of the interstitial space surrounding tissues. “Micro” refers to the extremely small scale and “dialysis” refers to the movement of chemicals across a semipermeable membrane. The microdialysis technique relies on the passive diffusion of substances across a dialysis membrane driven by a concentration gradient. The microdialysis probe is intracerebrally implanted in the target site, and a physiological salt solution (perfusate) is slowly and constantly pumped through the probe’s semipermeable membrane and is equilibrated with the surrounding tissue fluid. Intracerebral microdialysis can target ventricular CSF^4^, produced by the choroid plexus, which bathes the space between the arachnoid and pia mater. CSF is probably the most informative obtainable fluid for neurodegenerative disease prognosis, diagnosis and monitoring^5^. For instance, CSF has more physical contact with the brain compared to blood, as it is not separated from the brain by the tightly regulated blood-brain barrier^5^. As a result, proteins or peptides that may be directly reflective of neuropathology would be more likely to diffuse into CSF than any other bodily fluid. One of the most sensitive biomarkers for diagnosing AD are CSF concentrations of amyloid-β (Aβ) and tau^4^.

AD is the most common dementia among the elderly^6^. Two key neuropathological hallmarks that characterize AD are extracellular Aβ plaques and intracellular neurofibrillary tangles (NFTs), which are composed of aggregated Aβ peptides and hyperphosphorylated tau protein, respectively. Aβ (molecular weight ∼40-200 kDa^7^) refers to peptides that are released into circulation following the proteolysis of the transmembrane protein APP^8^. The most common peptides generated from APP cleavage are Aβ40 and Aβ42, with the latter isoform implicated in the formation of toxic oligomers, fibrils, and plaques in the diseased brain. Reduced levels of soluble Aβ42 in the CSF are associated with AD in patients and animal models^9^, leading to the adoption of this peptide as an AD biomarker. Tau proteins (molecular weight ∼55-62 kDa^10^) are produced by the alternative splicing from the gene *MAPT* and their primary roles are to maintain the stability of microtubules in axons. Increased levels of total tau (t-tau) in the CSF are associated with neurodegeneration, while the presence of phosphorylated tau (p-tau) in the CSF is associated with NFTs in AD patients and animal models^9^. It is believed that the decrease of Aβ42 in CSF reflects its aggregation and deposition in the brain parenchyma, whereas the increase in CSF tau reflects its extracellular release after neuronal degeneration and NFT formation^9^. Thus, these biomarkers are considered to be directly linked to the molecular pathogenesis of AD.

However, the feasibility of using microdialysis to sample large proteins (such as Aβ and tau) presents numerous challenges due to the loss of perfusion fluid through the high molecular weight cut-off dialysis membrane. This can be avoided by reducing the backpressure caused by the perfusion fluid coming from the probe inlet. We have achieved this in a mouse model of AD by performing microdialysis in a novel push-pull mode using a syringe pump to perfuse the microdialysis probe (push) and a peristaltic pump to collect the sample coming from the probe outlet (pull; Fig. 1). We have validated this method for longitudinal sampling over 12 months, and for the collection of two large proteins, along with two different isoforms of each protein, from the same microdialysis probe. Here we report for the first time how application of this technique allows for longitudinal measurement of Aβ and tau levels from CSF samples in the 3xTg AD mouse model^1^, prior to the development of pathology, and up until after both pathological Aβ and tau are present. Having validated the technique, we also provide specific recommendations for longitudinal sampling of large proteins *in vivo*. Moreover, the European directive for animal welfare in laboratory environments calls for the three R’s: replacement, reduction and refinement. Our method is an excellent example of the reduction and refinement principles, as it enables consecutive sampling from one animal without sacrifice, and with minimal disturbance to the animal’s mobility^11^. This novel method has many potential avenues; including translational research between preclinical models and patients, collecting large molecules from hard-to-reach tissues, and drug infusions aimed at ameliorating neuropathology.

**Figure 1.**
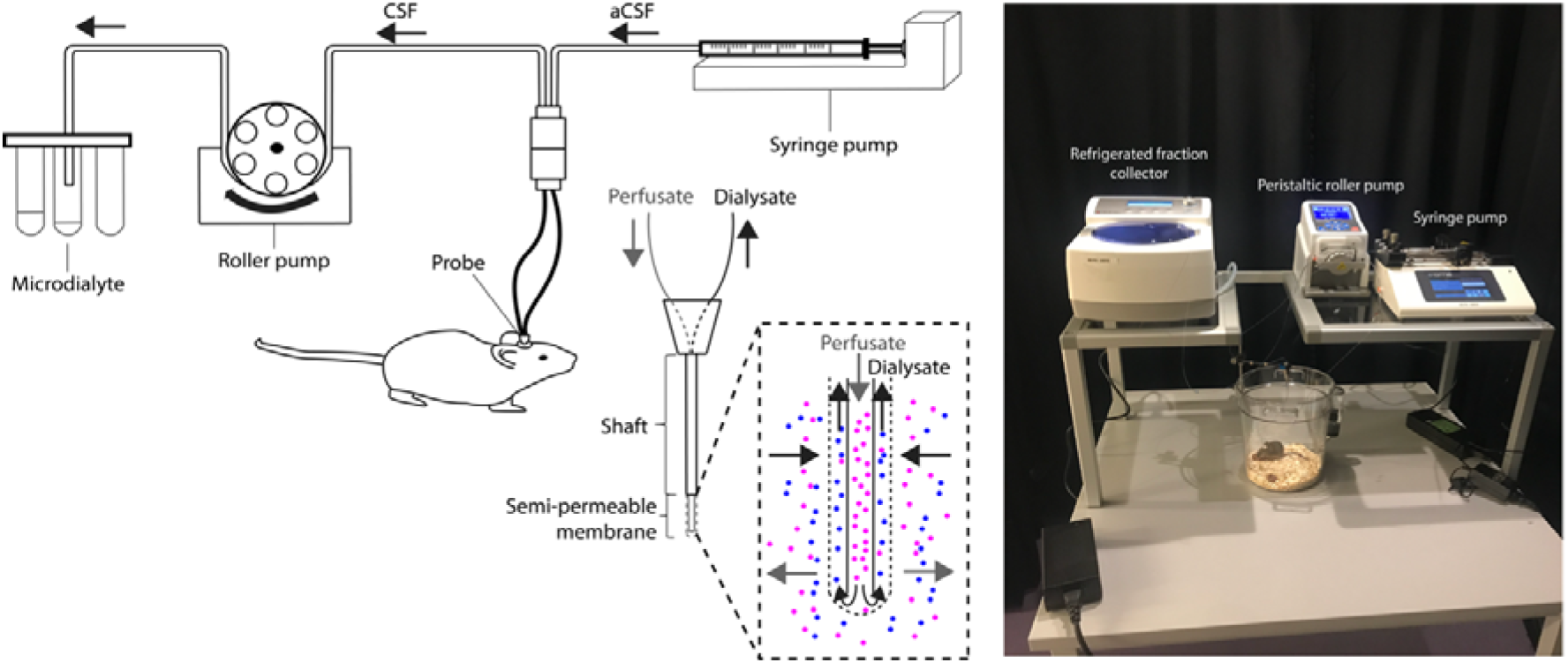
Push-pull microdialysis system and a probe with a high molecular weight cut-off membrane. The syringe pump is used to perfuse the microdialysis probe (push) and the peristaltic pump is used to collect the sample coming from the probe outlet (pull). The microdialyte is collected in a refrigerated fraction collector. The microdialysis probe consists of an outer semipermeable membrane with a specific pore size that allows for the recovery of large proteins. Figure adapted from Takeda and colleagues^12^.

## RESULTS

### *In vitro* testing of microdialysis probes and flow rates

We examined the influence of flow rate on recovery by comparing two flow rates: 1 and 0.1 µl/min (Fig. 2). The *in vitro* sampling using a microdialysis probe with a high molecular weight cut-off membrane allowed for the recovery of both Aβ and tau as measured by multiplex ELISA. However, the slower the flow rate, the higher the recovery of analytes, especially tau.

**Figure 2.**
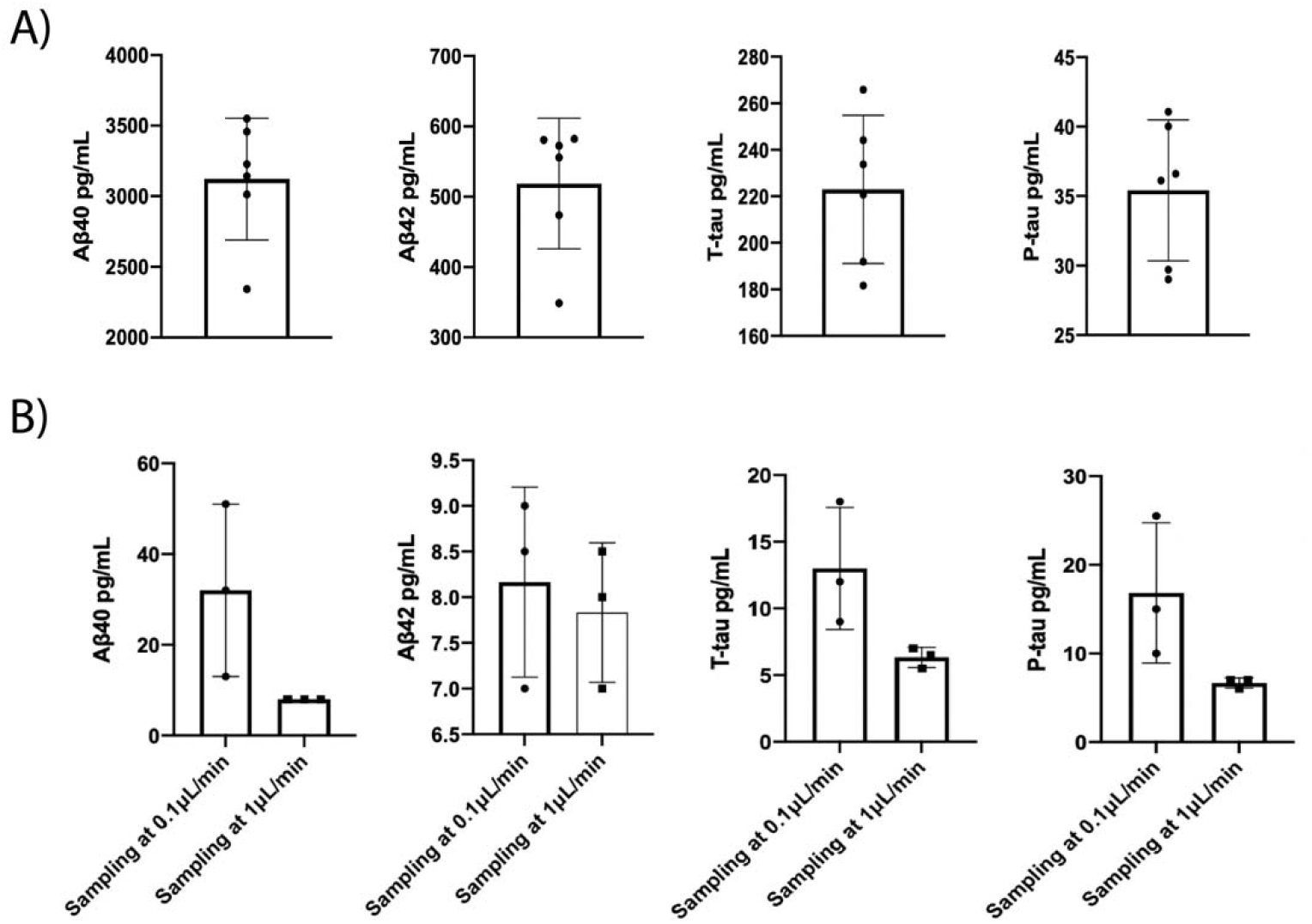
A) *In vitro* CSF samples from three healthy human subjects (Neurological Research Biobank, St. Olav’s Hospital, Trondheim, Norway) showing mean concentrations of Aβ40, Aβ42, t-tau and p-tau. Data displays duplicates of CSF samples from each experiment. B) We analyzed a non-microdialyte sample (and tested two flow rates: 1 and 0.1 µl/min) of CSF from three healthy human subjects using push-pull microdialysis followed by multiplex ELISA of Aβ40, Aβ42, t-tau and p-tau. Data is the mean from two duplicates of CSF samples. Error bars = SD.

One way to validate *in vitro* sampling techniques is to assess the percentage difference within protein recovery over multiple sampling sessions. Within the analytes, *in vitro* samples differed by 13.03 % for Aβ40, 8.77 % for Aβ42, 3.59 % for t-tau and 2.09 % for p-tau. The relative recovery (RR) for each analyte was calculated (Eqn. 1; Methods) based on the bulk concentration from lumbar puncture CSF samples from healthy human subjects (Neurological Research Biobank, St. Olav’s Hospital, Trondheim, Norway). The RR for Aβ40 was 5.14 %, for Aβ42 1.49 %, for t-tau 10.28 % and for p-tau 9.13 %.

### *In vivo* testing of microdialysis probes and flow rates

We examined CSF samples from three 3xTg AD mice and compared two flow rates (0.5 and 0.2 µl/min; Fig. 3). The *in vivo* sampling using a microdialysis probe with a high molecular weight cut-off membrane (Fig. 4) enabled the recovery of both Aβ and tau as measured by multiplex ELISA. However, the slower the flow rate, the higher the recovery of molecules, especially for Aβ42. Sampling at 0.2 µl/min was an adequate rate for recovering all four proteins of interest (Fig. 3), therefore all samples henceforth were sampled at this rate. There was an increase in the concentrations of CSF Aβ and tau from 3 to 3.5 months of age (m) in 3xTg AD mice (Fig. 5).

**Figure 3.**
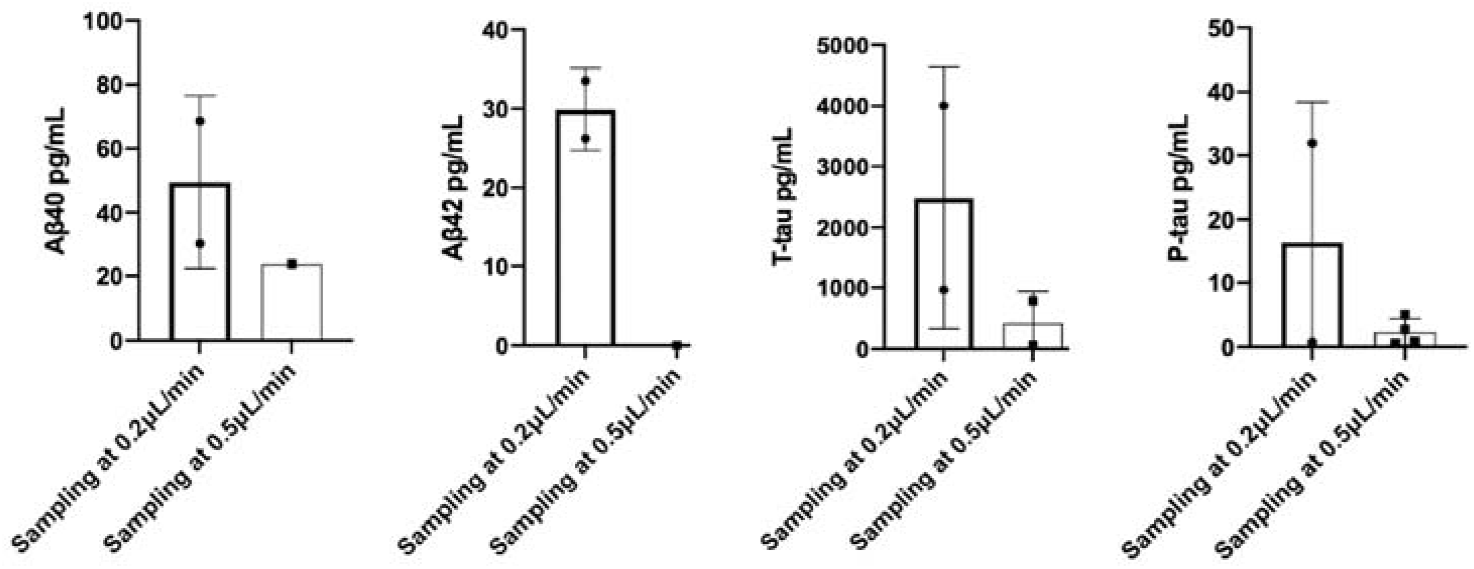
*In vivo* CSF microdialysis results from four 3xTg AD mice between 3 and 6 m showing mean concentrations of duplicates of Aβ40, Aβ42, t-tau and p-tau CSF samples as measured by multiplex ELISA. We examined two flow rates (0.2 and 0.5 µl/min) over eight experiments. No data points = no recovery of protein in microdialyte.

**Figure 4.**
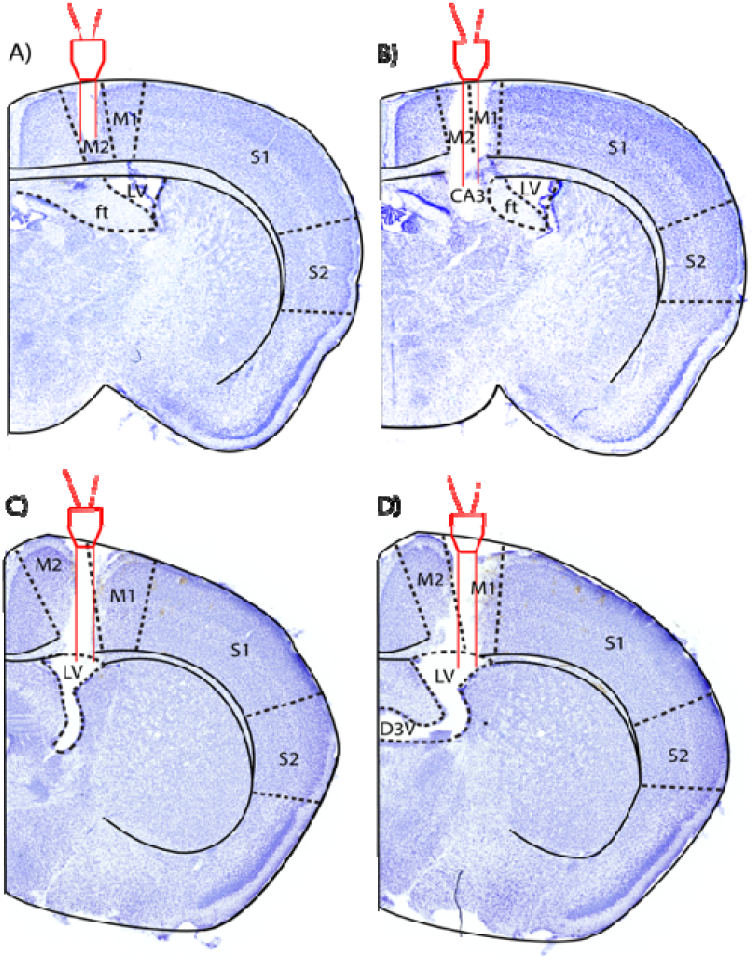
Microdialysis probe implantation location as verified with Cresyl violet (Nissl) staining in four 3xTg AD mice. A-B) The microdialysis probe was implanted posteriorly in the rostrocaudal axis to LV. The microdialysis probe was implanted in M2 and CA3. Stereotaxic coordinates: A/P −1 mm, M/L +1 mm, D/V −3 mm. C-D) The microdialysis probe was successfully implanted in LV along the rostrocaudal axis. Stereotaxic coordinates: A/P: - 0.1 mm, M/L: +1.2 mm, D/V −2.75 mm. Delineations based on Paxinos and Franklin^13^. Abbreviations; M2: secondary motor cortex; M1: primary motor cortex; S1: primary somatosensory cortex; S2; secondary somatosensory cortex; LV: lateral ventricle; ft: fimbria of the hippocampus; D3V: dorsal third ventricle; CA3: field CA3 of the hippocampus.

**Figure 5.**
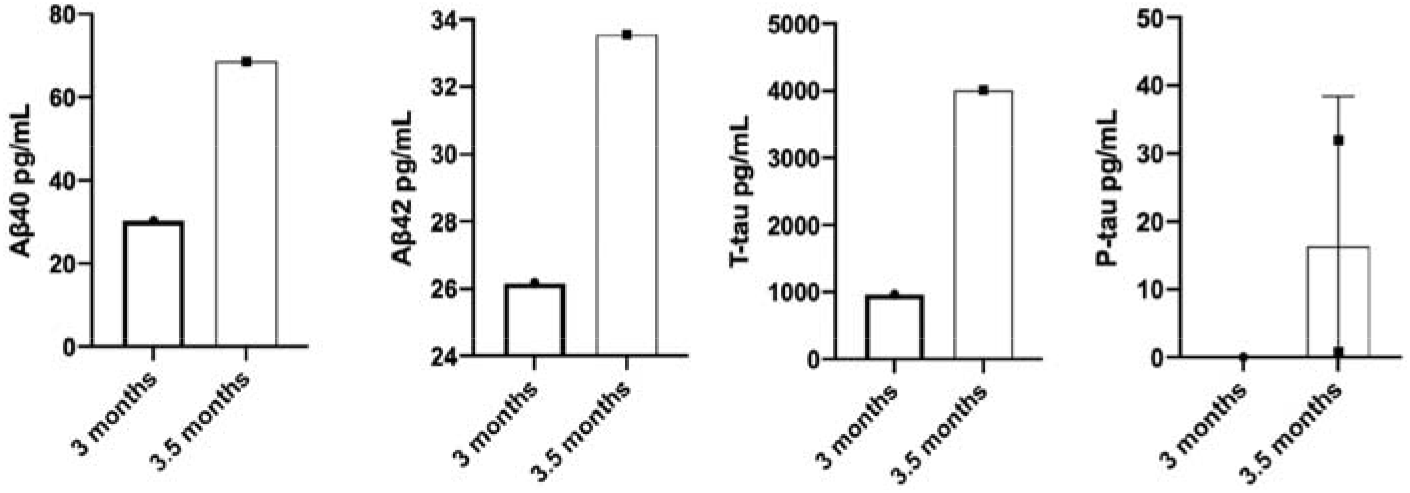
*In vivo* CSF microdialysis results from four 3xTg AD mice at 3 and 3.5 m showing mean concentrations of duplicates of Aβ40, Aβ42, t-tau and p-tau CSF samples as measured by multiplex ELISA. Data are from four experiments. No data points = no recovery of protein in microdialyte.

### Longitudinal *in vivo* microdialysis sampling of CSF

Longitudinal microdialysis sampling of sixteen animals from 1 until 18 m revealed a gradual increase in both CSF Aβ40 and Aβ42 concentrations, until 8-10 m when there was a decrease in CSF Aβ, followed by a slight increase and a plateau in concentrations as measured by multiplex ELISA (Fig. 6; see Methods for more details). The age at which CSF Aβ concentrations decreased coincides with when this transgenic mouse line develops Aβ plaques in the brain parenchyma (Fig. 7). The amount of t-tau and p-tau appears to be more variable between age groups until 10 to 11 m, when there is an increase in CSF tau concentrations until 18 m (Fig. 6). Higher levels of tau proteins are observed once NFTs are deposited in the brain of mice at 10-11 m (Fig. 8). There is a significant inverse correlation between CSF Aβ40/Aβ42 concentrations after Aβ plaques are deposited in the brain at 9 m and 18 m (Aβ40; *r*(167) = *-1*.*0, p* < 0.001, Fig. 9a, 10a; Aβ42: *r*(167) = *-0*.*999, p* = 0.001, Fig. 9b, 10b). Meanwhile, there is a significant positive correlation between CSF t-tau/p-tau at 11-12 m and when NFTs are deposited in the brain at 18 m (t-tau: *r*(167) = *0*.*976, p* = 0.02, Fig. 9c, 10c; p-tau: *r*(167) = *1, p* < 0.01, Fig. 9d, 10d). See Methods for more details regarding calculations of correlations and linear regression analyses.

**Figure 6.**
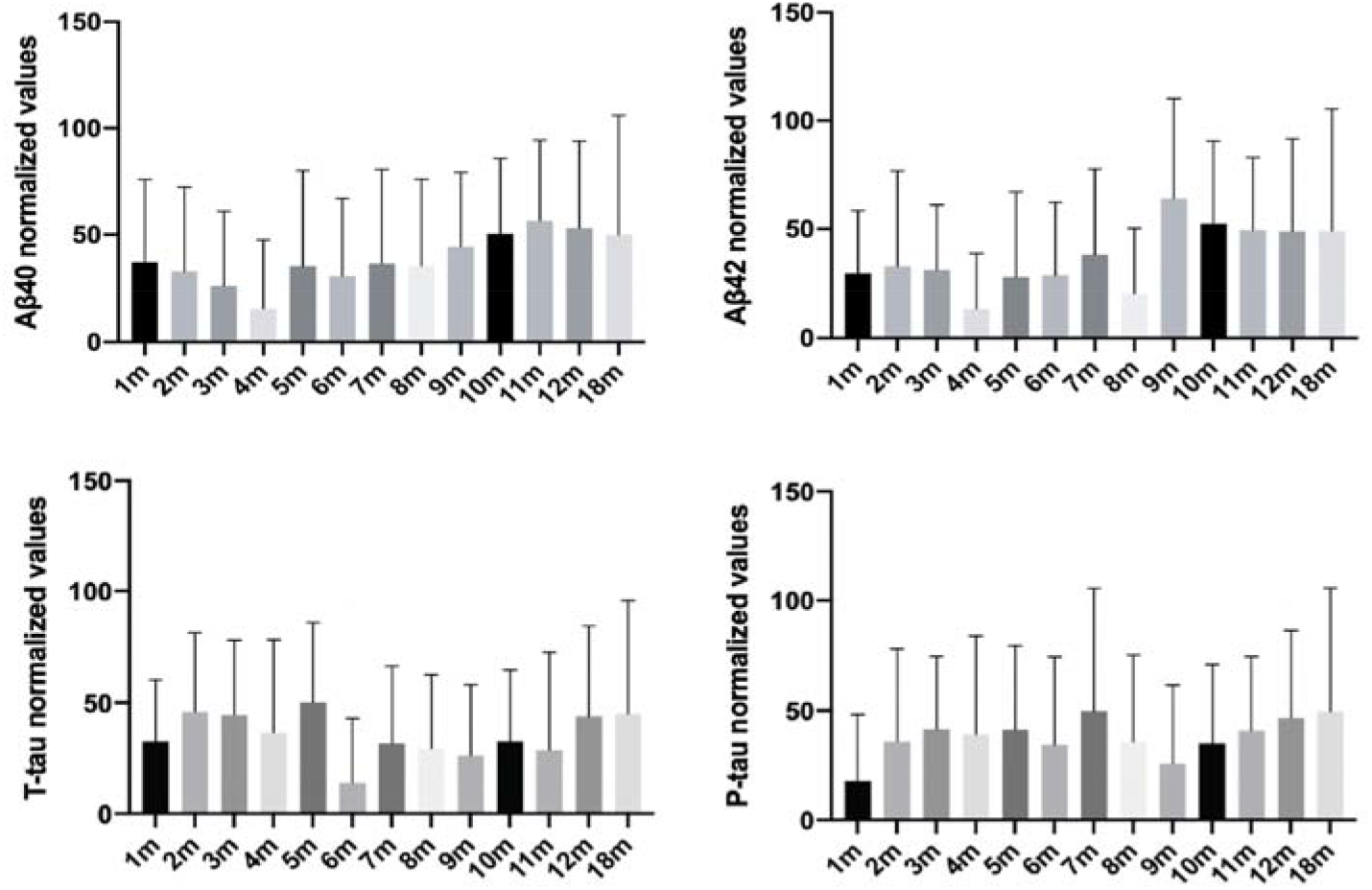
Longitudinal *in vivo* CSF microdialysis results from sixteen 3xTg AD mice, showing normalized concentrations in pg/ml of Aβ40 and Aβ42, t-tau and p-tau duplicates of CSF samples at different ages. Error bars = SD.

**Figure 7.**
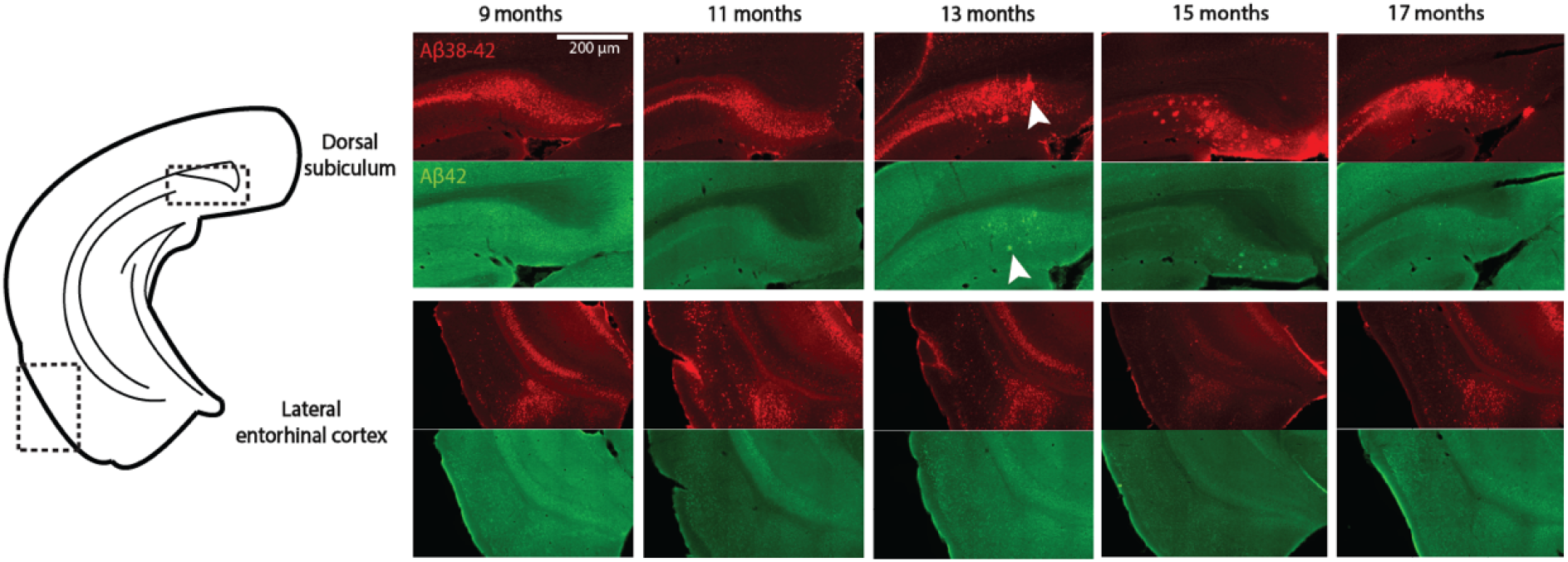
Aβ plaque deposition in five 3xTg AD mice at different ages. Aβ plaques are first apparent in the dorsal subiculum at 13 m in this transgenic mouse line (arrow heads). Red: Aβ38-42 (McSA1 antibody); green: Aβ42 (IBL Aβ42 antibody).

**Figure 8.**
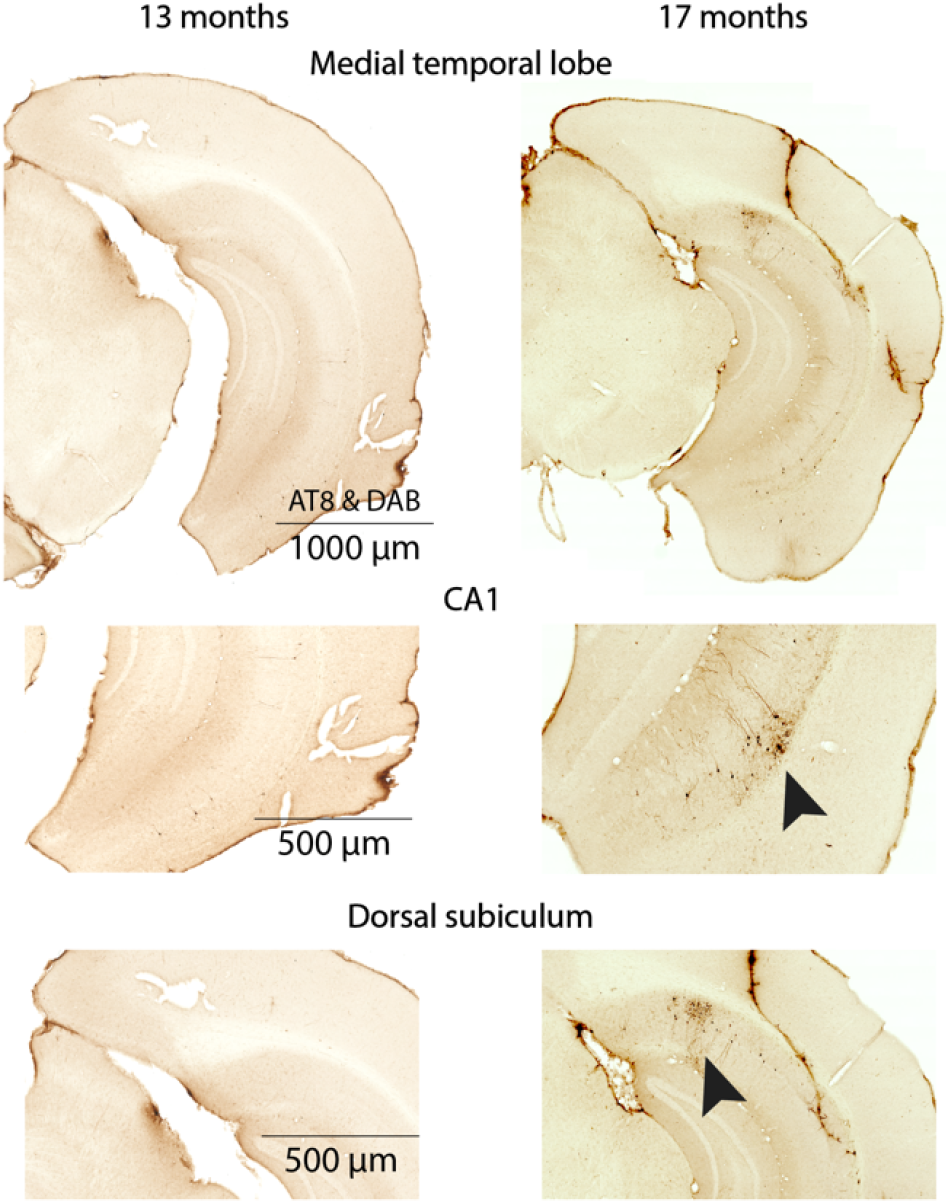
NFT deposition in two 3xTg AD mice at different ages. NFTs are first apparent in CA1 and the dorsal subiculum at 17 m in this transgenic mouse line (arrow heads). NFTs are labelled by the AT8 antibody and visualized with DAB.

**Figure 9.**
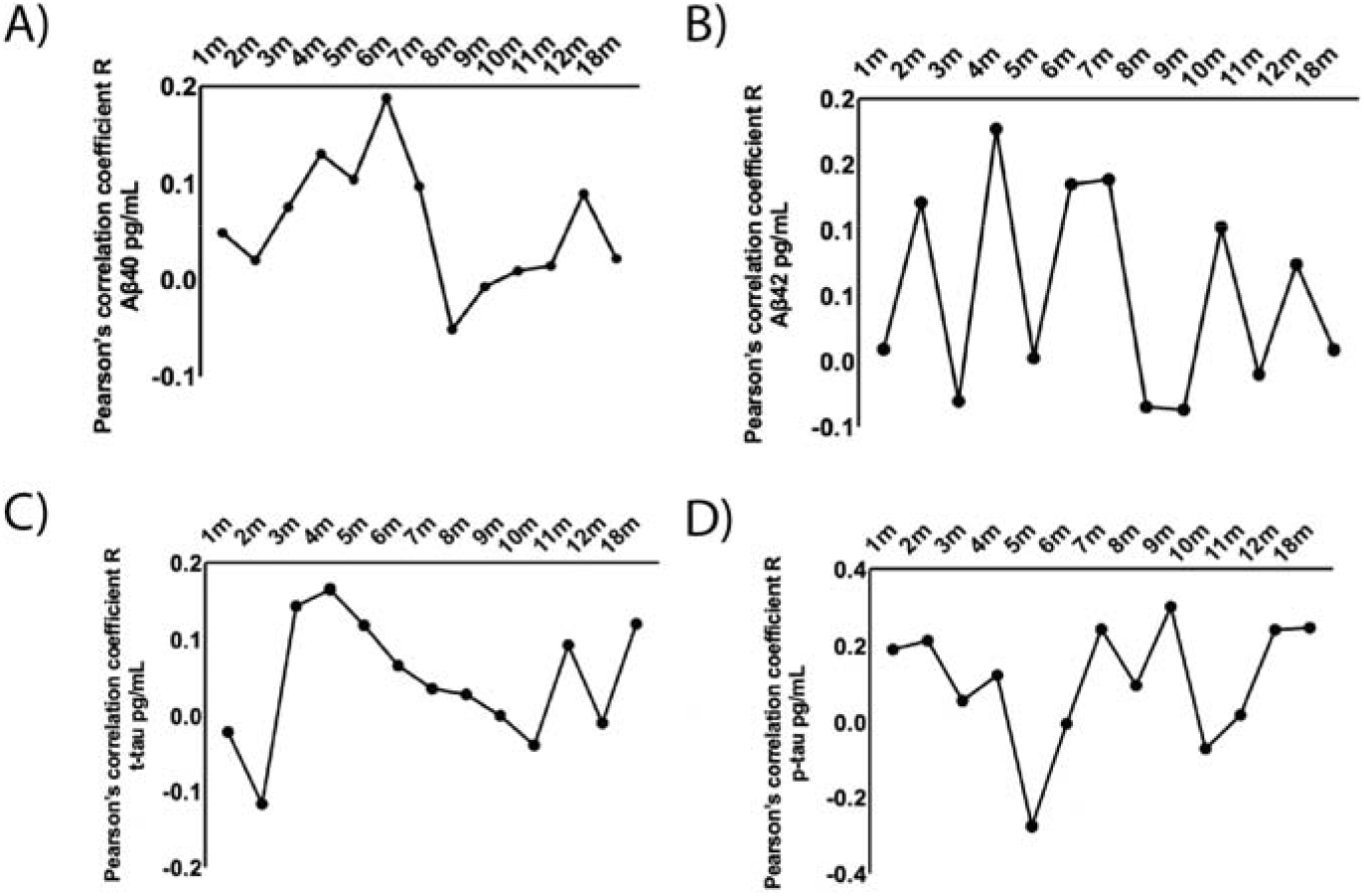
Pearson correlational scatterplots for A) Aβ40, B) Aβ42, C) t-tau, and D) p-tau, in sixteen 3xTg AD mice. Values represent Pearson’s correlation coefficient R of CSF protein concentrations at different ages in sixteen 3xTg AD mice.

**Figure 10.**
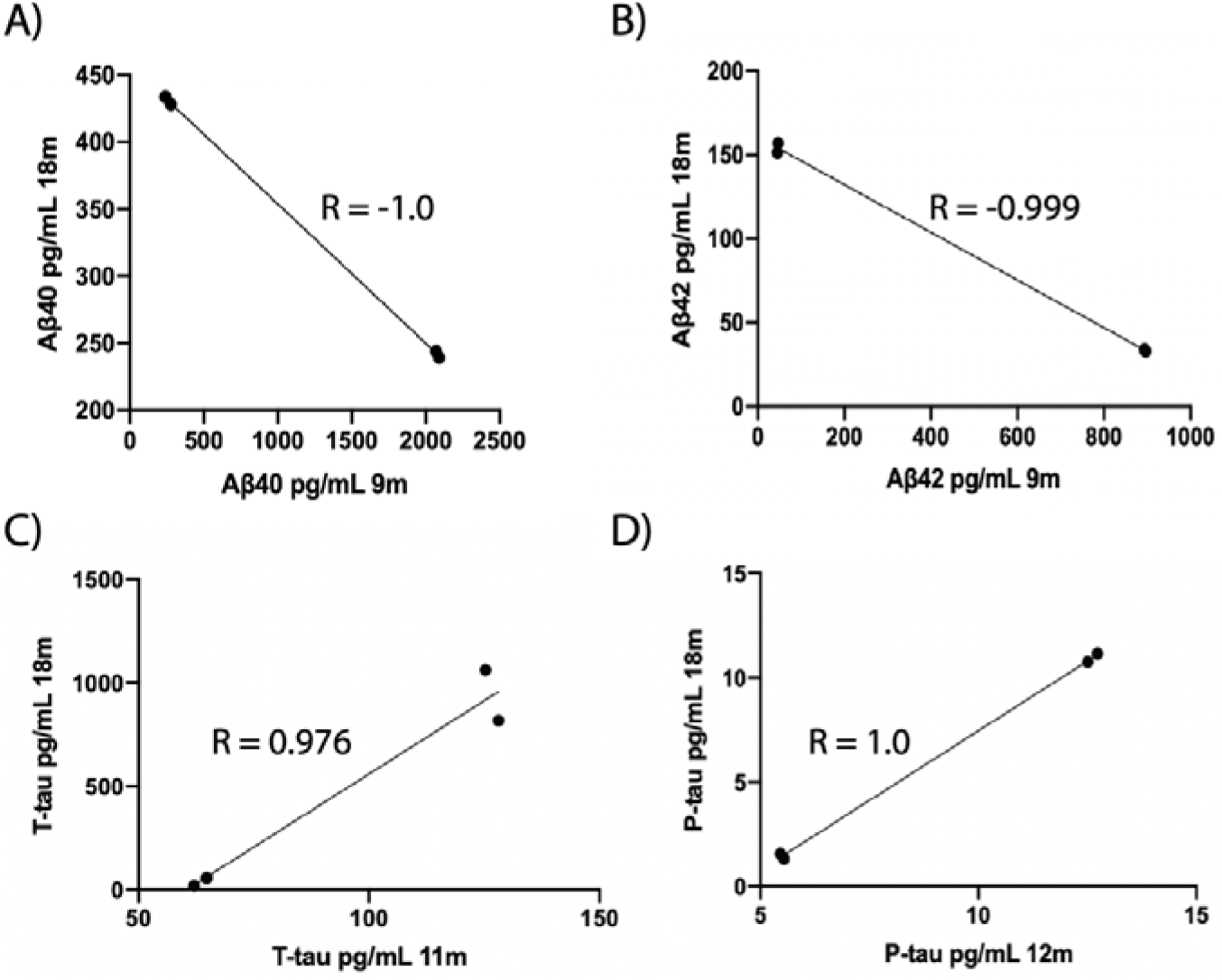
Correlational scatterplots for A) Aβ40 in 9 and 18 m, B) Aβ42 in 9 and 18 m, C) t-tau in 11 and 18 m, D) p-tau in 12 and 18 m from sixteen 3xTg AD mice. Lines represent linear regression displaying Pearson’s correlation coefficient R.

## DISCUSSION

Our novel push-pull microdialysis method enables longitudinal, simultaneous sampling of two large proteins, namely Aβ and tau and their phosphorylated forms. We hereby demonstrate not only that we can successfully follow the pathophysiological alterations in relevant CSF biomarkers along the AD disease cascade in a preclinical model, but also that these changes reflect equivalent ones observed in AD patients during disease progression. The application of our novel microdialysis technique for longitudinal CSF sampling thus further proves the translational validity of the 3xTg AD mouse model in the study of progressive AD pathology. Furthermore, successful application of longitudinal microdialysis can provide valuable insights into disease onset and staging, as well as aid the development and assessment of the efficacy of relevant treatments for AD neuropathology^4^. In contrast to classical CSF sampling in animals, such as drainage from the cisterna magna after sacrificing the animal^4^, our microdialysis method allows consecutive sampling from the same animal over time and causes minimal disturbance to the animal. As a result, we can use animals as their own intrinsic controls, as done in the clinic with patients, and as such fully utilize these valuable experimental subjects by maximizing research data and relevant findings from preclinical models.

We found that the slower the flow rate, the better the recovery of analytes in the microdialyte. For recovery of tau proteins, slower flow rates are pertinent. Although our novel method allows for the recovery of Aβ40, Aβ42, p-tau and t-tau, the recovery rates for Aβ are low, with slightly higher recovery for tau proteins. This could be due to the low migration rate of Aβ across the dialysis membrane or post-sampling factors, such as centrifugation, temperature and blood in the CSF sample^14^. The higher recovery for tau may be due to the protein being more stable post-sampling. Generally, the low recovery of large proteins may be due to the amount of time used to sample the microdialyte, as it has been found that the recovery of large proteins to decrease significantly after a few hours^15^. However, sampling from animals that model a progressive neurodegenerative disease is novel, and is thus of great value in the study of intra-variability of proteins that aggregate in AD. Importantly, our findings suggest that longitudinal CSF Aβ sampling in transgenic mice mirrors what is observed in AD patients and also correlates with what is observed with positron emission tomography (PET) scans^4,16^; there is a gradual increase in both Aβ isoforms, with a gradual decrease in Aβ concentrations once Aβ plaques are deposited into the brain parenchyma followed by a slight increase and plateau in Aβ levels. Meanwhile, CSF tau levels are more variable early in the disease progression but appear to steadily increase after NFTs are deposited in the brain, as is observed in patients.

The relative recovery of an analyte is inversely proportional to the perfusate flow rate^17-21^. For instance, at extremely slow flow rates (<0.1 μl/min) it is possible to reach 100 % recovery of an analyte^22-24^, but this recovery also depends on both the diffusional characteristics of the analyte of interest, as well as dialysis membrane type and length. Using rates of 0.057 μl/min^24^ and 0.1 μl/min^22^, researchers have obtained close to 100 % recovery of their analytes of interest. Based on these findings, it is possible to reduce the flow rate to achieve near equilibrium between the actual concentration of the analyte and the concentration in the microdialyte. One also needs to keep in mind that the slower the flow rate, the longer the animal needs to stay in the sampling session, and this should be limited as much as possible without compromising the recovery rate.

Optimization of the push-pull microdialysis setup requires specific tools: i) A reliable syringe pump system (Fig. 1) for the perfusate is needed for accurate microdialysis sampling. ii) The perfusate microsyringe or injector (Fig. 1) needs to maintain accurate, constant flow rate as incorrect or varying flow rates will alter recovery or exchange of substances across the microdialysis membrane^25^. iii) The peristaltic pump (Fig. 1), which sets a flow rate identical to that for the syringe pump to avoid backpressure in the sampling apparatus. iv) A programmable refrigerated fraction collector (Fig. 1) can facilitate the microdialysis sampling procedure by circumventing the risk of evaporation of microdialytes. v) A microdialysis probe (Fig. 1) consists of an outer semi-permeable membrane with specific pore sizes and which has a wide variety of molecular weight cut-offs. The diffusion rate across the microdialysis membrane is dependent on several factors including concentration gradient and molecular size of the protein, probe membrane surface area and temperature. Studies have shown that a longer membrane yields better recovery of proteins^17,18,26-28^. Larger proteins that would be able to pass through the semi-permeable membrane pores may have a low migration rate across the membrane.

Perfusate composition is important because its components diffuse into the tissue in the same way that the analyte of interest diffuses into the microdialysis probe. Ideally, the perfusate should contain the exact concentration of all solutes that diffuse through the probe membrane and that are found in the extracellular fluid, except of course for the protein of interest. As solutes are removed from the external medium by the probe, a region surrounding the probe may become depleted of solute^25^ and affect the dialysis of the protein of interest. Therefore, an equilibration period is required before sample collection can start. The time to reach intracerebral equilibrium in rodents is most commonly 7 days^29,30^, but there might be conditions that demand longer equilibration periods. The insertion of the microdialysis probe in the brain is thought to induce only minor damage to the tissue, however, some bleeding might occur at the insertion point as there is cerebral vasculature in close proximity to the lateral ventricle^31^.

Researchers have examined the influence of trauma on microdialysis measures^32-36^, and found that neurotransmitter release sites adjacent to the microdialysis probe were damaged. They further proposed that the trauma caused by the microdialysis probe suppressed neurotransmitter uptake, and therefore their microdialytes underestimated true extracellular concentration. Moreover, traumatic brain injury (TBI) has been found to be a risk factor for developing AD^37^, and results in an increase in CSF S100B protein levels^38^. In line with this, while using intracerebral microdialysis in patients, researchers found that extracellular concentrations of S100B change markedly in response to brain injury after TBI; therefore, this protein could be an elegant biomarker for neuronal injury.

We found that glial inflammation does not necessarily increase in the brain after chronic guide implantation, but it cannot be excluded that the ventricle could dilate, and that cortical atrophy might be observed in some animals. However, enlarged ventricles and concomitant brain atrophy could be a possible indicator of AD disease progression^39^. In line with this, a study has shown that in the 3xTg AD mouse model, ventricles become enlarged compared to control mice already at 2 m^40^. Moreover, we observe an increase in CSF p-tau levels in transgenic mice as they age, and this protein is the main component of paired helical filaments which form NFTs and which are involved in the formation of extracellular senile plaques^41^. Therefore, CSF p-tau levels may be considered as a potential biomarker for AD, in addition to the standardized use of CSF t-tau levels^4^, as p-tau directly represents axonal degeneration. With axonal degeneration, the survival of neurons would be compromised and lead to atrophy, as well as ventricular enlargement, secondary to brain matter loss. Activation of the brain’s macrophages and other immune cells may exacerbate both Aβ and tau pathology, and is today considered a core neuropathological hallmark in AD^42^. Based on our results, however, we have no indication that chronic microdialysis guide implantation causes unexpected inflammation and can therefore exclude this factor as contributing to exacerbated neuropathology (Supplementary Fig. 1). This is in line with our findings of a decrease in CSF Aβ concentrations once amyloid plaques are present (Fig. 8), and an increase in CSF tau once NFTs are deposited (Fig. 9).

### Future perspectives and conclusions

Direct comparison between microdialysis studies is not straightforward because the procedure may differ between study protocols and the microdialysis guide cannula can be implanted in different ways^4^. The analysis method used to ascertain concentrations of analytes may vary between laboratories and differ in sensitivity. For instance, the multiplex ELISA kit used in the current study may display some cross-reactivity between human and murine CSF samples, as the kit is reactive to human Aβ and tau. Only 25 µl of CSF are needed for our analysis, which is a significant advantage when collecting samples from a small animal. Our study shows for the first-time longitudinal CSF sampling in animals modelling AD. Previous studies on CSF protein concentrations from lumbar puncture have shown that at approximately 5 m there should be 1500 pg/mL of Aβ^43^, and at 3 m one should observe 15,000 ng/ml of tau^44^ in mice. One would expect lower concentrations of Aβ and tau in ventricular CSF samples because lumbar puncture in mice is commonly a terminal procedure without the possibility of serial sampling, and larger volumes of CSF are collected^4^. Moreover, ventricular CSF samples collected by microdialysis may become diluted, and the concentrations of Aβ and tau have been shown to vary between the ventricles and lumbar regions of animals and patients^4^. Meanwhile, in the APP/PS1 transgenic mouse line, CSF t-tau concentrations were approximately 500 pg/ml in young animals by collections from the cisterna magna^45^. The 3xTg AD transgenic line used in our study harbours a mutant human tau transgene (*MAPT*), and therefore tau levels may be higher, but sampling from the cisterna magna is another terminal procedure with larger volumes of CSF collected. Therefore, future research may find it applicable to compare CSF protein concentrations from lumbar puncture or drainage from cisterna magna with microdialytes.

Conducting microdialysis sampling in freely moving animals, as demonstrated in our study, can provide many data points for a single animal, obtained under the same experimental paradigm, and thereby reveal the relationship between concentrations of analytes and time. Although intracerebral microdialysis has mainly been applied to animals, previous studies have demonstrated that this method might be a valuable tool for the monitoring of patients during neurointensive care^46,47^. For instance, Marklund and colleagues^48^ used intracerebral microdialysis catheters with a high molecular cut-off membrane to harvest interstitial t-tau and Aβ42 proteins in patients with TBI, and found that both proteins constituted sensitive biomarkers for axonal injury, which, as mentioned, is an important risk factor for developing AD. Future studies using longitudinal intracerebral microdialysis for analysing AD progression may thus also include sampling of additional biomarkers, such as S100B proteins. In terms of translatability between transgenic mice and patients, mice and humans share a large amount of corresponding genes, as almost every gene found in mice or humans has been observed in a closely related form in the other^4^. Therefore, there is a pressing need and also a unique opportunity for better quality data from preclinical animal models investigating biomarkers, which can be translated to patients. The core biomarkers for diagnosis in AD patients (Aβ42, t-tau, and p-tau) have been found to translate well across species, whereas biomarkers of inflammation translate to a lesser extent between mouse models and patients^4^. Towards genuine translation and improved understanding of biomarker changes as a result of AD pathology progression, it is imperative to longitudinally follow preclinical models, akin to clinical follow up, in order to elucidate key pathological mechanisms of the disease and also to determine which of these models ultimately carry substantial validity to help treat patients.

## ACKNOWLEDGEMENTS

This study was supported by the Liaison Committee for Education, Research and Innovation in Central Norway (Samarbeidsorganet HMN-NTNU) grant number 2018/42794 and the Joint Research Committee (Felles Forskningsutvalg - St. Olav’s Hospital HF and the Faculty of Medicine, NTNU) grant number 2019/26157. CSF samples collected by lumbar puncture from healthy human subjects were generously donated by Professor Emerita Linda White, Department of Neuromedicine and Movement Science, NTNU, Trondheim from the Neurological Research Biobank (St. Olav’s Hospital, Trondheim, Norway).

## AUTHOR CONTRIBUTIONS

Conceptualization, C.B, A.S. and I.S.; Methodology, C.B.; Investigation, C.B., C.L. and M.H.; Writing – Original Draft, C.B.; Writing – Review & Editing, C.B., I.S., A.S., M.H., T.H.F. and C.L.; Funding Acquisition, C.B., A.S. and T.H.F.; Supervision, A.S. and I.S.

## ONLINE METHODS

### Recovery of analytes and *in vitro* validation

Recovery of molecules reflects the concentration in the dialysate of the molecules of interest in relation to the true concentration surrounding the microdialysis membrane^29^. A high recovery means that the concentration in the microdialyte is close to the true concentration in the CSF of a specific molecule (or protein) of interest. Recovery in microdialysis is dependent on many factors, such as the perfusion flow rate, diffusion rate (of the protein and of the probe membrane), and the performance of the probe (cut-off, diameter and length of the membrane, chemical interaction between proteins and the membrane). Therefore, a crucial parameter in microdialysis is RR^49-51^, which entails the extraction efficiency used to determine the true analyte concentration in the extracellular fluid (*in vivo*) or in a bulk sample (*in vitro*). RR can be calculated as the concentration of the analyte in the dialysate (C_d_) divided by the concentration in the bulk sample (C_b_), multiplied by 100 (Eqn. 1).

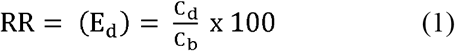

Probe *in vitro* recovery tests should be conducted prior to actual experiments to assess the efficiency and dialyzing properties of the microdialysis probe for the particular protein of interest. The purpose of these experiments is to determine if the flow rate and collection period selected yield a sample with a detectable concentration of analyte, as measured by recovery. In addition to the chosen probe, here we also used a syringe pump for fluid perfusion (CMA 4004; CMA Microdialysis AB, Kista, Sweden), an *in vitro* storage stand (CMA 130; CMA Microdialysis AB, Kista, Sweden), probe clips (CMA Microdialysis AB, Kista, Sweden), and a peristaltic pump (Reglo ICC digital; CMA Microdialysis AB, Kista, Sweden) for *in vitro* recovery testing. It is however important to note that *in vitro* RR estimates have been previously used to correct *in vivo* microdialyte values, but findings from other studies suggest that this method is imprecise^52^, since diffusion in tissue differs significantly from diffusion in a solution^53^. In order to reduce backpressure and potential leakage of the system, addition of vent holes to the outlet of the microdialysis probe has been reported by researchers^54^, this is however inapplicable in our system as the peristaltic pump reduces the systemic pressure.

At the beginning of our experiments, all microdialysis probes were perfused with artificial CSF at a flow rate of 60 μL/min in order to fill the entire system with the perfusate. A 7-day equilibration period was always allowed before starting *in vivo* CSF sampling. All samples were collected in polypropylene plastic vials (CMA Microdialysis AB, Kista, Sweden) that were sealed with plastic caps for storage (as recommended by the multiplex enzyme-linked immunosorbent assay [ELISA] kit used for analyses). During the *in vitro* calibration the effect of time and flow rate on recovery was examined. A microdialysis probe was placed in CSF collected from healthy subjects (Neurological Research Biobank, St. Olav’s Hospital, Trondheim, Norway) and perfused at 0.1 and 1 μL/min (see results section 5.1). Based on these studies, the flow rate was changed to 0.2 μL/min and a constant sampling volume of 60 μL was collected. All samples were stored at −80 °C until further analyses.

### Patient CSF samples

Three CSF samples collected by lumbar puncture from healthy subjects (Neurological Research Biobank, St. Olav’s Hospital, Trondheim, Norway), with written consent for the use in research (REK 2013/467, Neurological Research Biobank) were used for *in vitro* experiments.

### Animals

A total of twenty-eight 3xTg AD mice^55^ were used in this study. The 3xTg AD mouse line, which harbors the Swedish *APP*, the P301L *MAPT*, and M146V *PSEN1* mutations, was generated by Jackson laboratories and obtained from MMRRC (strain name: B6;129-Tg(APPSwe,tauP301L)1Lfa*Psen1tm1Mpm*/Mmjax). Four 3xTg AD mice were used to refine lateral ventricle implantation surgeries, seven were used for characterization of neuropathology, and the remaining mice were implanted at 1-month-of-age, except for one animal that was implanted at 6 m, and two animals that were implanted at 8 m. A total of two B6129 mice were used in this study. Mouse with identical genetic backgrounds to the 3xTg AD mouse line, generated by Jackson laboratories, were used as age-matched controls (strain name: B6129SF2/J). The two B6129 mice were implanted at 1-month-of-age. All mice were housed individually following surgery in diurnal light conditions (12-hour light/dark cycle) and had *ad libitum* access to food and water.

Animal living conditions and breeding were approved by the Norwegian Animal Research Authority and were in accordance with the Norwegian Animal Welfare Act §§ 1-28, the Norwegian Regulations of Animal Research §§ 1-26, and the European Convention for the Protection of Vertebrate Animals used for Experimental and Other Scientific Purposes (FOTS numbers 17392/21061).

### Pilot experiments of microdialysis guide implantation surgeries

Four 3xTg AD mice were used to refine microdialysis probe implantation surgery. In two of these mice, the probe was implanted in the CA3 field of the hippocampus, and in two animals, the probe was implanted in the lateral ventricle (Fig. 4). In the first two animals, the probe was implanted more posteriorly to the lateral ventricle in the rostrocaudal axis, and in the animals that were implanted in the lateral ventricle, CSF surfaced shortly after the insertion of the guide cannula during implantation surgeries. There are arteries in close proximity of the lateral ventricle that can be inadvertently hit, and minor bleeding was observed in a few animals after implanting the microdialysis guide (see Discussion). This needs to be avoided, because the blood could clog the microdialysis probe and make it difficult to sample CSF from the animal.

### Microdialysis guide cannula implantation surgery

Implantation surgery was performed to insert microdialysis guide cannulas (CMA 7; CMA Microdialysis AB, Kista, Sweden) into the lateral ventricle of mice. Mice were anesthetized with isoflurane gas (4 % induction and 1.5-3 % maintenance; IsoFlo vet., Abbott Laboratories, Chicago, IL, USA) prior to being fixed in a stereotaxic frame (Kopf Instruments; Chicago, IL, USA). Anaesthesia levels were monitored throughout surgery via respiration and testing toe pinch reflexes. All surgical instruments were autoclaved prior to surgery and disinfected during surgery in 75 % ethanol. After being placed into the anaesthesia mask, the animal was fixed into the stereotaxic frame using ear bars while placed prone on a heating pad. The fur of the skull was shaved, and the skin was cleaned with saline and chlorhexidine. Prior to making any incisions, Marcain (0.03-0.18 mg/kg; Aspen Pharma, Ballerup, Denmark) was injected subcutaneously into the scalp and Metacam (5 mg/kg; Boehringer Ingelheim Vetmedia, Copenhagen, Denmark) and Temgesic (0.05-0.1 mg/kg; Invidor UK, Slough, Great Britain) were administered subcutaneously for intraoperative pain relief. After using bregma and lambda to horizontally level the skull, the stereotaxic coordinates were derived to target the lateral ventricle (A/P −0.1 mm, M/L +1.2 mm, D/V - 2.75 mm; Fig. 11). The microdialysis guide cannula was attached to the stereotaxic frame using a guide clip and connection rod for the clip (CMA Microdialysis AB, Kista, Sweden). The skull was drilled through at these coordinates and the guide cannula was slowly lowered into the drilled hole. The guide cannula was attached to the skull with super glue and dental cement (Dentalon Plus; Cliniclands AB, Trelleborg, Sweden). Following surgery, the animal was placed in a heated chamber until awake and moving normally. Post-surgery, Metacam and Temgesic were administered within 24 hours.

**Figure 11.**
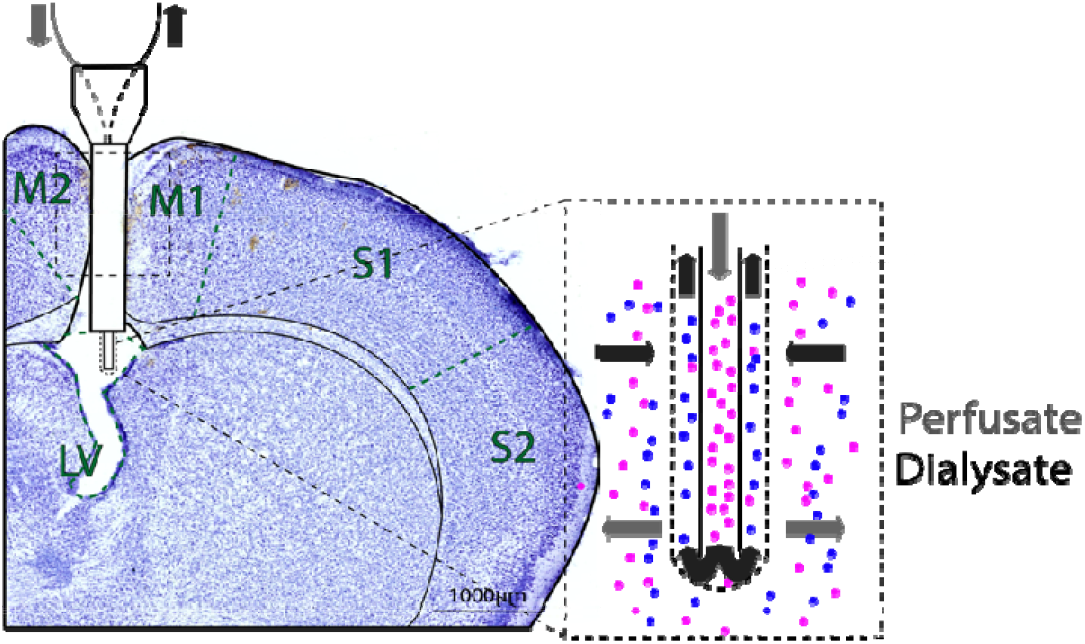
Microdialysis guide cannula implanted into the LV of the mouse brain. The microdialysis probe with a high molecular weight cut-off membrane replaces the dummy probe during microdialysis sampling. Abbreviations; M2: secondary motor cortex; M1: primary motor cortex; S1: primary somatosensory cortex; S2: secondary somatosensory cortex; LV: lateral ventricle.

### Push-pull microdialysis apparatus and sampling

Intracerebral microdialysis was performed for the dynamic sampling of CSF Aβ and tau. The push-pull method was utilized to allow for the measurement of larger endogenous proteins with a high molecular weight. A refrigerated fraction collector (CMA 470) was set to 6 °C for the storage of collected CSF in polypropylene plastic vials. Fluorinated ethylene propylene (FEP) peristaltic tubing (CMA Microdialysis AB, Kista, Sweden) was placed inside each plastic vial for collection and connected to the cassette of the peristaltic roller pump (Reglo ICC Digital). This peristaltic FEP tubing was connected to the outlet side of the microdialysis probes (β-irrigated 2 mDa microdialysis probe; CMA 7; CMA Microdialysis AB, Kista, Sweden) with a polyethersulfone 2 mm membrane with tubing adapters bathed in 75 % ethanol. FEP tubing (CMA Microdialysis AB, Kista, Sweden) was connected to each microsyringe. The FEP tubing was then connected to the inlet part of the microdialysis probes. Transparent cages were prepared with 1.5 cm of bedding, filled water bottles, and treats. Artificial CSF (CMA Microdialysis AB, Kista, Sweden) that consists of NaCl (147 mmol/l), KCl (2.7 mmol/l), CaCl_2_ (1.2 mmol/l) and MgCl_2_ (0.85 mmol/l) was used. This artificial CSF was mixed with 4 % bovine serum albumin (BSA; 30 % in saline; Sigma-Aldrich, Saint-Louis, MO, USA) to allow for better recovery of Aβ and tau^54^, and to minimize the chance of microdialyte fluid loss^56^. This solution was made fresh prior to each sampling session. Artificial CSF was loaded inside a gastight microsyringe (CMA Microdialysis AB, Kista, Sweden), which was placed into a syringe pump (CMA 4004). The ‘dead volume’ of the FEP outlet tubing (1.2 μL/100mm) was calculated. 100 cm of FEP outlet tubing was used, and therefore the first 12 μL sampled from each animal was discarded. Mice were habituated to human handling and restraining for one week prior to microdialysis sampling and restrained in order to remove the dummy probe of the guide cannula and replace it with the microdialysis probe. Prior to inserting the microdialysis probes into the guide cannula, the probe was conditioned in 75 % ethanol for better recovery of analytes^54^. After the mice were connected to the sampling apparatus, the room lights were turned off for the duration of the experiment. The animals were awake and freely moving while connected to the sampling apparatus to avoid any artifacts produced by anesthesia. At the conclusion of microdialyte sampling, the vials of 60 µl CSF were centrifuged and kept at −80 °C until the samples were analyzed with multiplex ELISA. All tubing was discarded after one sampling session, as the 4 % BSA clogged the tubing.

### Quantification of Aβ and tau concentrations in CSF

The MILLIPLEX® MAP human amyloid beta and tau magnetic bead panel 4-plex kit (Millipore, Burlington, MA, USA) and the Bio-Plex 200 System instrument (Biorad, Hercules, CA, USA) were used to assess simultaneously the concentrations of Aβ40, Aβ42, tTau, and pTau (Thr181) in CSF samples. The samples were undiluted and analysed in duplicates.

### Tissue processing and immunohistochemistry

Mice were administered a lethal dose of sodium pentobarbital and transcardially perfused with Ringer’s solution followed by paraformaldehyde (PFA, 4 %) in 125 mM phosphate buffer (PB). Brains were extracted and fixed for a minimum of 24 hours in PFA at 4 °C and transferred to a 2 % dimethyl sulfoxide solution prepared in PB for 24 hours at 4 °C. Brains were sectioned coronally at 40 µm on a freezing microtome.

Immunohistochemical processing was conducted on tissue from chronically implanted animals, as well as non-implanted transgenic mice for characterization of AD neuropathology. During fluorescent immunohistochemistry, heat-induced antigen retrieval (HIAR) was carried out on all tissue at 60 °C for 2 hours in PB. Then, sections were washed 3 times for 10 minutes with PB containing 0.2 % Triton X-100 (Merck, Darmstadt, Germany; PBT). Next, sections were blocked using 5 % normal goat serum (Abcam, Cambridge, UK) in PBT (PBT+) for 1 hour before incubation with the primary antibody. Sections were incubated with the primary antibodies Iba1 (1:1000; ab15690, Abcam, Cambridge, UK), McSA1 (1:1000; MediMabs, Montreal, Canada; characterization), and Aβ42 (1:1000; IBL America, Minnesota, USA; characterization) in PBT+ for 4 hours at 4 °C. In order to label and visualize primary antibodies, directly conjugated Alexa Fluor secondary antibodies (Invitrogen, California, USA) at a 1:400 concentration was used. First, sections were washed 3 times for 10 minutes with PBT. Then, sections were incubated with Alexa Fluor 546 to visualize Iba1 and McSA1, and Alexa Fluor 488 to visualize Aβ42, for 2 hours at room temperature, protected from light. Then sections were washed for 10 minutes with 4 ′, 6-diamidino-2-fenylindol (DAPI; 1:10 000; Sigma-Aldrich, Saint-Louis, MO, USA) and PB, followed by 3 washes for 10 minutes with PB.

In order to characterize NFT deposition, tissue from non-implanted transgenic mice was stained with a free□floating procedure using the monoclonal antibody AT8 (ThermoFisher, Massachusetts, USA). HIAR was carried out on all tissue at 60 °C for 2 hours in PB. After washing with PB (2 × 10 minutes) and permeabilizing with 0.5 % Triton□X□100 in Tris□buffered saline (TBS□Tx; 50 mm Tris, 150 mm NaCl, pH 8.0) for 10 minutes, the tissue was blocked with 10 % goat serum in TBS□Tx for 30 minutes and incubated with the primary antibody (AT8, 1:1000) in TBS□Tx overnight at 4 °C. The following day, the sections were washed with TBS□Tx (3 × 10 minutes), incubated with a biotinylated goat anti□mouse secondary antibody (1:500; Sigma□Aldrich, St Louis, MO, USA) in TBS□Tx for 90 minutes before washing with TBS□Tx (3 × 10 minutes) and incubating with ABC (Vectastain ABC kit, Vector Laboratories, Burlingame, CA, USA) for 90 minutes. After washing with TBS□Tx (3 × 10 minutes) and Tris□HCl (50 mm Tris, pH 7.6, 2 × 5 minutes), tissue was incubated with 0.67 % diaminobenzidine (DAB) and 0.024 % H2O2 for 10 minutes before a final wash with Tris□HCl (2 × 5 minutes).

Processed tissue was mounted on cut edge frosted glass slides (VWR International, Radnor, PA, USA) from a solution of 50 mM tris(hydroxymethyl)aminomethane (Merck Millipore, Merck kGaA, Darmstadt, Germany; pH 7.6) and then left to dry for at least 4 hours on a 38 °C heating plate, protected from light. Mounted tissue was placed in Xylene (VWR International, Radnor, PA, USA) for at least 5 minutes for defatting and to remove excess water from the tissue, and then coverslipped with Entellan (VWR International, Radnor, PA, USA) containing Xylene. The mounted tissue was then left to dry overnight, protected from light.

One series of each brain were dehydrated in ethanol, cleared in xylene and rehydrated before staining with Cresyl violet (Nissl; 1g/l) for 3 minutes. The sections were then alternatively dipped in ethanol–acetic acid (5 ml acetic acid in 1l 70 % ethanol) and rinsed with cold water until the desired differentiation was obtained, then dehydrated, cleared in xylene and coverslipped with Entellan containing Xylene.

Sections were scanned using a Mirax-midi scanner (objective 20X, NA 0.8; Carl Zeiss Microscopy, Oberkochen, Germany), using either reflected fluorescence (for sections stained with a fluorophore) or transmitted white light (for sections stained with Cresyl Violet; Nissl) as the light source.

### Statistics

No statistical methods were used to predetermine sample sizes, but our sample size is similar to those reported in previous publications^24^. In order to normalize data from CSF sample concentrations of Aβ40, Aβ42, t-tau, and p-tau (Fig. 6), the mean of duplicates from sixteen 3xTg AD mice were assigned a value between 0 and 1 in order to visualize trends between different age groups. The correlation between CSF protein concentrations and age (Fig. 9) were based on Pearson’s correlation coefficient R and corresponding p-values from the raw data displayed as normalized values (Fig. 6). The correlations between Aβ and tau at respective ages (Fig. 10) were based off correlations between CSF protein concentrations and age (Fig. 9) and visualized to further highlight the strong relationship between protein deposition in the brain and protein CSF concentrations. A simple linear regression model was also applied to this data (Fig. 10). All statistical tests and graphs were made in Prism 9 (GraphPad Software Inc., CA, USA).

## SUPPLEMENTARY FIGURES

**Supplementary figure 1.**
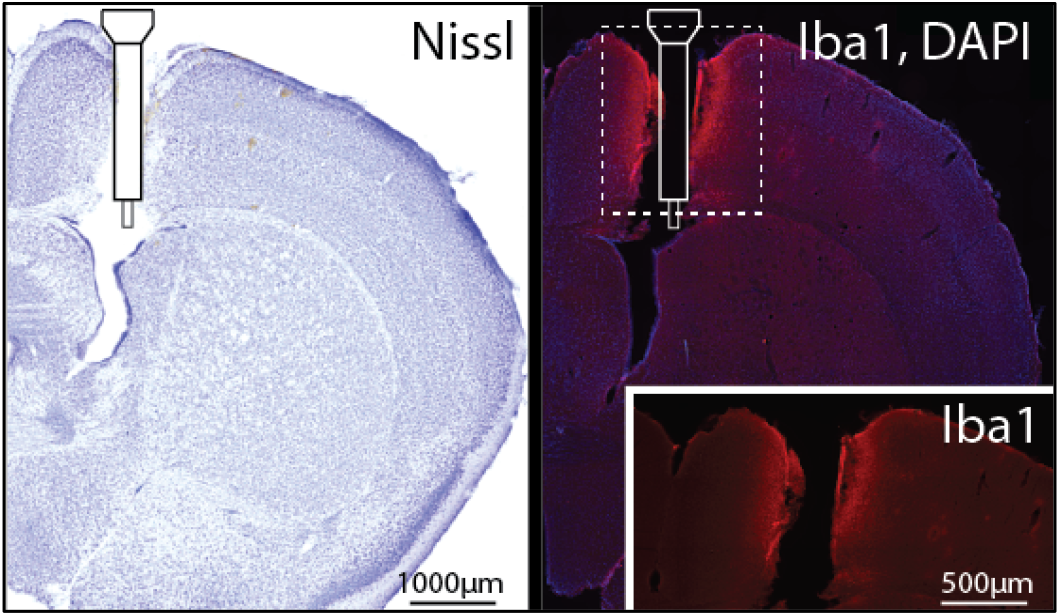
Microdialysis guide cannula implantation in the lateral ventricle and inflammation. Cresyl violet (Nissl) stained section showing the implantation track. Iba1 (glial inflammation; red) and DAPI (nucleic staining; magenta) of the same section reveals minor neuroinflammation along the implanted track.

